# Simultaneous Triple-Parametric Optical Mapping of Transmembrane Potential, Intracellular Calcium and NADH for Cardiac Physiology Assessment

**DOI:** 10.1101/2021.09.09.459667

**Authors:** Sharon A George, Zexu Lin, Igor R Efimov

**Author notes:** **Contact Information:** SAG ZL.

## Abstract

Investigation of the complex relationships and dependencies of multiple cellular processes that govern cardiac physiology and pathophysiology requires simultaneous dynamic assessment of multiple parameters. In this study, we introduce triple-parametric optical mapping to simultaneously image metabolism, electrical excitation, and calcium signaling from the same field of view and demonstrate its application in the field of drug testing and cardiovascular research. We applied this metabolism-excitation-contraction coupling (MECC) methodology to test the effects of blebbistatin, 4-aminopyridine and verapamil on cardiac physiology. While blebbistatin and 4-aminopyridine alter multiple aspects of cardiac function suggesting off-target effects, the effects of verapamil were on-target and it altered only one of ten tested parameters. Triple-parametric optical mapping was also applied during ischemia and reperfusion; and we identified that metabolic changes precede the effects of ischemia on cardiac electrophysiology.

## Introduction

Cardiac physiology is profoundly complex, and the careful coordination of numerous cell signaling pathways govern every single heartbeat. These physiological processes that vary beat to beat and long term are usually sub-divided into metabolic, electrical, and mechanical components governed by metabolism-excitation-contraction coupling (MECC). Electrical activity in the heart includes a sequence of opening and closing of ion channels and pumps which cause cardiomyocyte depolarization and repolarization. The resulting changes in the transmembrane potential (V_m_) in the cardiomyocytes are recorded as action potentials using electrical or optical methods. The electrical excitation of the heart serves as the trigger of a mechanical contraction. Excitation-contraction coupling or the translation of electrical excitation to mechanical contraction is controlled by cytosolic calcium (Ca^2+^) ion concentration, which increases following electrical excitation and Ca^2+^ ion binds to a contractile protein in the cardiomyocyte, triggering contraction. Both the electrical and mechanical processes of the heart require energy, which is provided in the form of ATP generated by metabolic processes in the mitochondria. Thus, all three components of cardiac function are intertwined into MECC. As such, studying this complex MECC phenomenon requires the ability to simultaneously assess these three facets of cardiac function. In this study we report a new approach to simultaneously image V_m_, Ca^2+^ and NADH (metabolic marker).

Optical mapping is a methodology that optically records cardiac physiology with high spatial and temporal resolution, either as autofluorescence of endogenous biological substances or as fluorescence of specifically designed dyes. Optical mapping of the heart was first applied to record V_m_ and then NADH autofluorescence^1,2^. Since then, optical mapping has also been applied to record Ca^2+ 3^. Dual parameter optical mapping of V_m_ and NADH as well as V_m_ and Ca^2+^ have also been applied in assessing cardiac physiology^4,5^. However, as described above, the three interdependent facets of cardiac function will all need to be measured simultaneously to develop a complete picture of cardiac physiological modulation by drugs or disease. In this study, we report for the first time, triple-parametric optical mapping for the simultaneous measurement of NADH, V_m_ and Ca^2+^.

Preclinical safety and efficacy testing are a crucial component of the drug development process. This step is important in determining dosing and toxicity which could include identifying off-target effects of drugs before clinical trials and approval for use in patients. Cardiotoxicity is the primary cause (19%) of drug withdrawal from the market in the United States^6^ and the second leading cause worldwide^6,7^, underscoring the need for efficient and thorough cardiac methodologies of drug screening. Current preclinical drug testing primarily focuses on the effects of drugs on the electrical activity (QT interval) and contractility of the heart^6,8^. While this is an essential first step, it does not give a complete picture of cardiac physiology modulation by drugs as it does not consider calcium handling or metabolic states of the cardiac tissue. Unexpected off target effects of the drugs being tested could result in serious complications or fatality. In this study, we present a novel approach to measure ten different important aspects of cardiac MECC using triple parametric optical mapping. We investigated the effects of three different compounds (blebbistatin, 4-aminopyridine and verapamil) using triple parametric optical mapping and present the data in a ten-parameter panel (TPP). TPP graphs include information on action potential upstroke, duration and conduction, intracellular calcium release and reuptake as well as the metabolic state of the heart.

Triple parametric optical mapping and TPP graphs could also benefit the study of complex cardiac diseases such as ischemia and reperfusion. Acute ischemic bouts are known to have multiple and severe effects on cardiac physiology. Ischemia has been previously demonstrated to alter the electrical activity^9–11^, calcium handling^12^ as well as the metabolic^2^ functions of the heart. However, the sequence and the interrelationship between these three aspects of cardiac MECC have not been studied simultaneously before, due to lack of appropriate methodology. In this study, we also determined the simultaneous modulation of multiple aspects of cardiac physiology by ischemia and their restoration during reperfusion. Thus, applying triple parametric optical mapping to study MECC during disease progression could provide valuable new targets for therapy.

## Results

Triple parametric optical mapping system was 3D printed and set up as illustrated in Figure 1. All design files for 3D printed hardware (STL format) and data analysis software (Matlab) are open source and available at Github (https://github.com/optocardiography).

**Figure 1.**
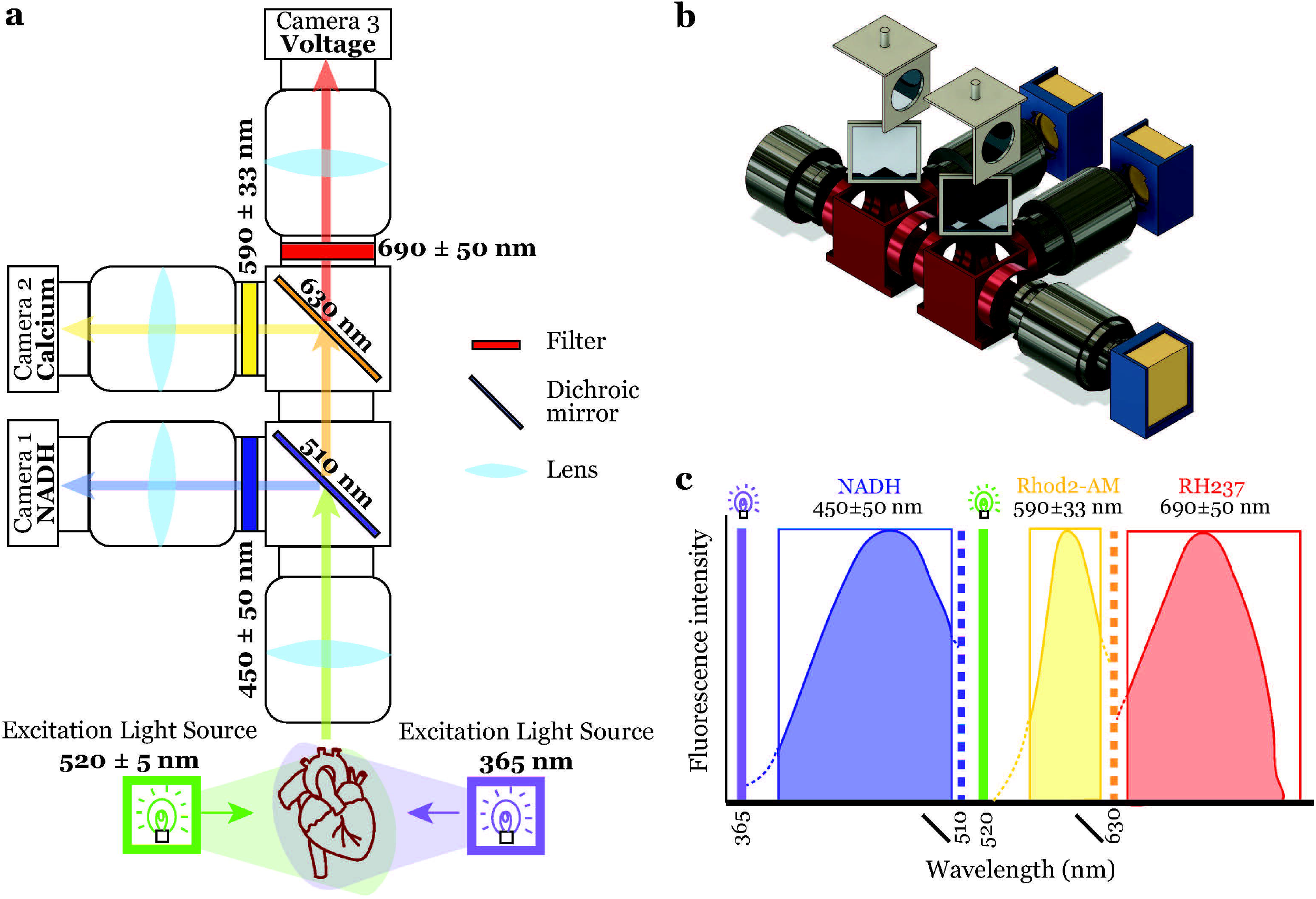
Triple parametric optical mapping system. **a)** Schematic of the triple parameter optical mapping system illustrating the optics used in the system and the light path for the three signals – V_m_, Ca^2+^ and NADH. **b)** 3D rendering of the triple parameter optical mapping system illustrating the 3D printed hardware that houses the optics used in the system. **c)** Spectra of the three parameter signals – NADH autofluorescence, Rhod2-AM (Ca) and RH237 (V_m_) fluorescence, illustrating the separation of the three signals as implemented in this system. Solid vertical lines: LED light source wavelength, dotted vertical lines: dichroic mirror, boxes: filters.

### Blebbistatin modulates cardiac physiology

Motion of the heart during image acquisition introduces artifacts in the recorded optical signals which hinder the analysis of repolarization-related parameters. To prevent these motion artifacts, electromechanical uncouplers such as blebbistatin are routinely used in optical mapping of V_m_ and Ca^2+^. Thus, the first step in this study was to analyze the effects of blebbistatin on the ten parameters of cardiac physiology.

Representative V_m_ and Ca^2+^ traces (Figure 2a top) illustrate motion artifacts that are introduced in the repolarization phases in Control (no treatment) hearts. Activation/intensity maps generated from these traces are shown in Figure 2b (top). Optical recordings during Control treatment allowed for the measurement of 7 parameters – V_m_ upstroke rise time (V_m_ RT), transverse and longitudinal conduction velocity (CV_T_ and CV_L_, respectively), anisotropic ratio (AR), Ca^2+^ upstroke rise time (Ca^2+^ RT), delay between V_m_ and Ca^2+^ activation (V_m_-Ca^2+^ delay) and NADH autofluorescence intensity (NADH). Restitution, which is the property of electrophysiological parameters to vary with diastolic interval (typically decrease with decreasing diastolic interval), was observed in CV_T_ and CV_L_.

**Figure 2.**
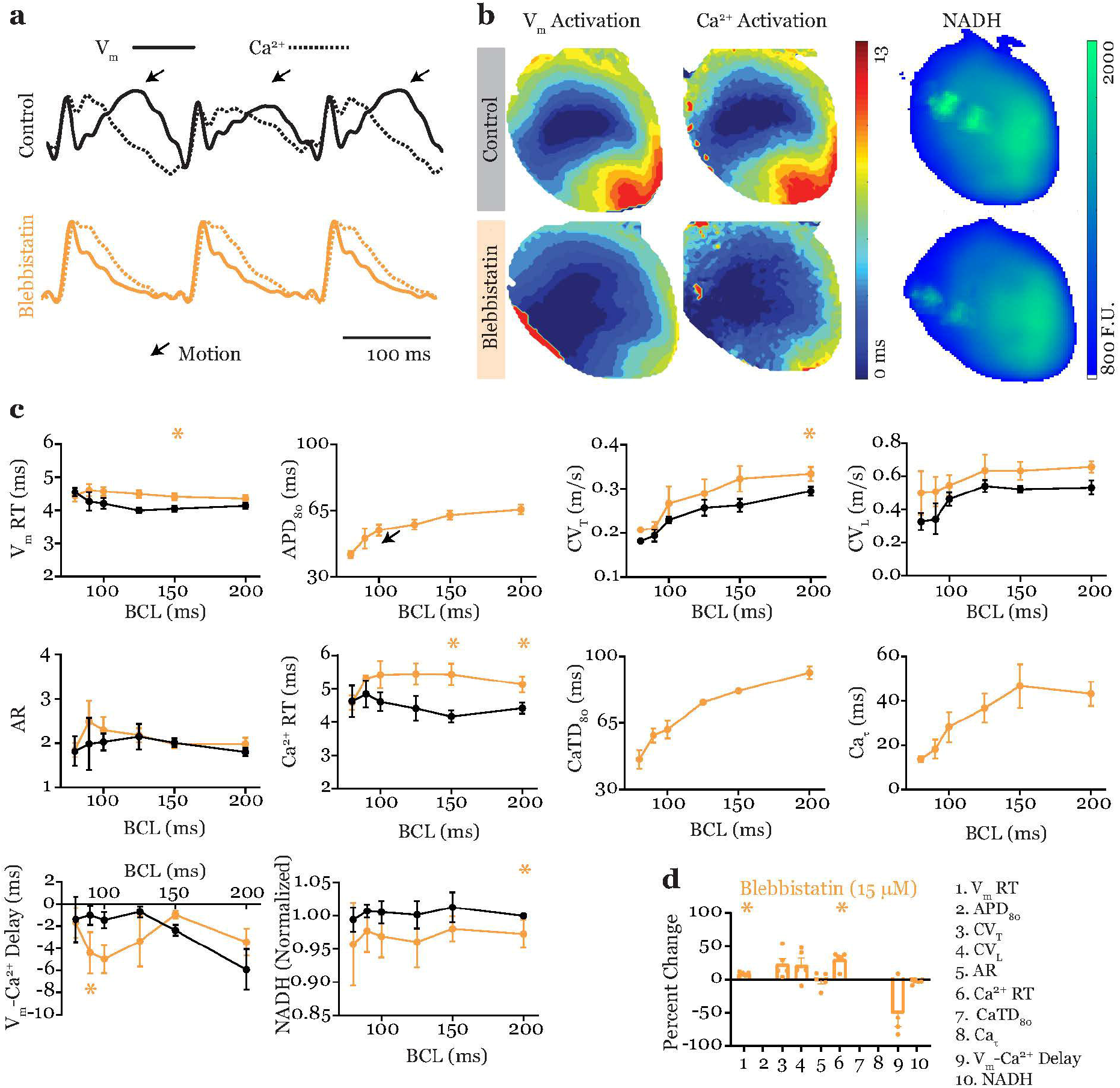
Electromechanical uncoupling in triple parametric optical mapping. **a)** Representative traces of V_m_ and Ca^2+^ (solid and dotted traces, respectively) recorded without (Control, black) and with the electromechanical uncoupler Blebbistatin (15 μM, orange). **b)** Representative V_m_ activation maps (left), Ca^2+^ activation maps (middle) and NADH intensity maps (right) recorded during Control and Blebbistatin treatment. **c)** Summary restitution properties of the ten parameters measured simultaneously by triple parametric optical mapping. **d)** Ten parameter panel showing the effects of Blebbistatin on cardiac physiology at 150 ms BCL, reported as percent change from Control. Data presented as mean ± s.e.m., n=5, * denotes p<0.05 versus Control using paired Student’s t-test with Bonferroni correction.

Treatment by blebbistatin (15 μM) abolished contractions and removed motion artifacts as shown in Figure 2a (bottom) and repolarization/calcium reuptake parameters such as action potential duration (APD_80_), calcium transient duration (CaTD_80_) and calcium decay rate constant (τ) were determined in addition to the 7 parameters measured during Control treatment. Restitution property was observed in APD_80_, CV_T_ and CV_L_. Additionally, CaTD_80_ and Ca_τ_ was also rate dependent. Specifically, both CaTD_80_ and Ca_τ_ decreased with increasing pacing rate (Figure 2c).

Blebbistatin also induced significant differences in cardiac physiology compared to Control. Blebbistatin increased both V_m_ and Ca^2+^ upstroke RTs as well as CV_T_. On the other hand, NADH intensity was reduced after blebbistatin treatment (Figure 2c). These effects were specific to slower pacing rates (BCL = 150, 200 ms) and differences versus Control were not significant at faster pacing rates. Finally, blebbistatin also increased V_m_-Ca^2+^ delay at 90 ms BCL.

In the TPP graph in Figure 2d, percent change in each of the ten parameters induced by blebbistatin with respect to control, at 150 ms BCL, is summarized which once again illustrates that blebbistatin significantly increases V_m_ and Ca^2+^ RTs.

### On- and Off-Target effects of Drugs on Cardiac Physiology

The application of triple parametric optical mapping in drug testing was then evaluated with two well studied drugs currently used in treating patients – 4-AP and verapamil. The effects of these drugs on cardiac physiology were tested in the presence of blebbistatin to be able to determine repolarization-/calcium reuptake-related parameters. Therefore, each physiological parameter during 4-AP and verapamil treatment were compared to blebbistatin treatment to determine significant drug-related effects. The effects of 4-AP and verapamil on cardiac physiology are summarized in Figure 3. The TPP graphs in Figure 3a, demonstrate the differences between 4-AP and verapamil. While 4-AP had multiple on-target and off-target effects on cardiac physiology, verapamil only had a specific on-target effect.

**Figure 3.**
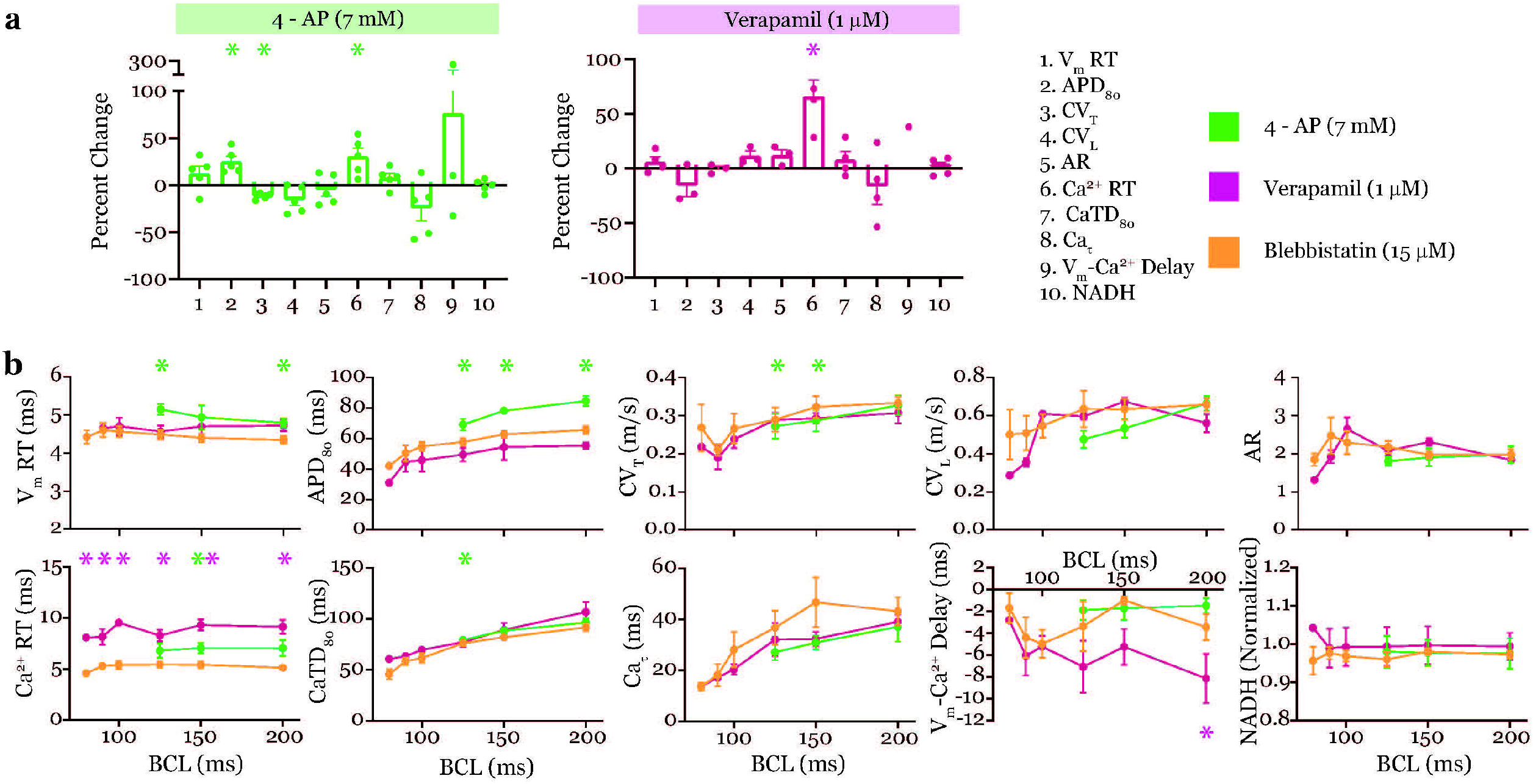
Triple parametric optical mapping in drug testing. **a)** Ten parameter panels demonstrating effects of 4-AP (7 mM, left, green) and Verapamil (1 μM, right, magenta). **b)** Summary restitution properties of the ten parameters measured simultaneously by triple parametric optical mapping. Data presented as mean ± s.e.m., n=5, * denotes p<0.05 versus Blebbistatin using paired Student’s t-test with Bonferroni correction.

#### Effects of 4-AP on Cardiac Physiology

Treatment with 4-AP (7 mM), a transient outward potassium current (I_to_) blocker, prolonged APD_80_ at all tested pacing rates compared to blebbistatin, as expected. 4-AP treatment also prevented pacing the hearts at pacing rates faster than 125 ms BCL. Additionally, 4-AP also prolonged Ca^2+^ RT and slowed CV_T_ at 150 ms BCL (Figure 3a). Furthermore, 4-AP also induced V_m_ RT and CaTD_80_ prolongation as indicated in the restitution graphs in Figure 3b.

#### Effects of Verapamil on Cardiac Physiology

In contrast to the multiple effects of 4-AP, the effects of verapamil, an L-type calcium channel blocker (I_CaL_) was specific to the upstroke of the calcium transient. Verapamil prolonged Ca^2+^ RT alone at all pacing rates tested as shown in Figure 3b.

### Acute modulation of Cardiac Physiology by No Flow Ischemia

A separate set of hearts were perfused with Control solution and a short episode of ischemia was induced by turning off the perfusion to the heart for a 5 min period followed by reperfusion. Ischemia modulated multiple parameters of cardiac physiology as shown in Figure 4 while reperfusion restored all of them to pre-ischemic (baseline) values. Activation/ intensity maps during baseline, ischemia (5 mins) and reperfusion (5 mins) are shown in Figure 4a. All three maps in each column were generated from optical data that was simultaneously recorded. Time-dependent responses of the 7 tested parameters during ischemia and reperfusion are illustrated in the graphs in Figure 4b while TPP graphs for ischemia and reperfusion demonstrating significant modulation of cardiac physiology during ischemia and restoration during reperfusion are shown in Figure 4c.

**Figure 4.**
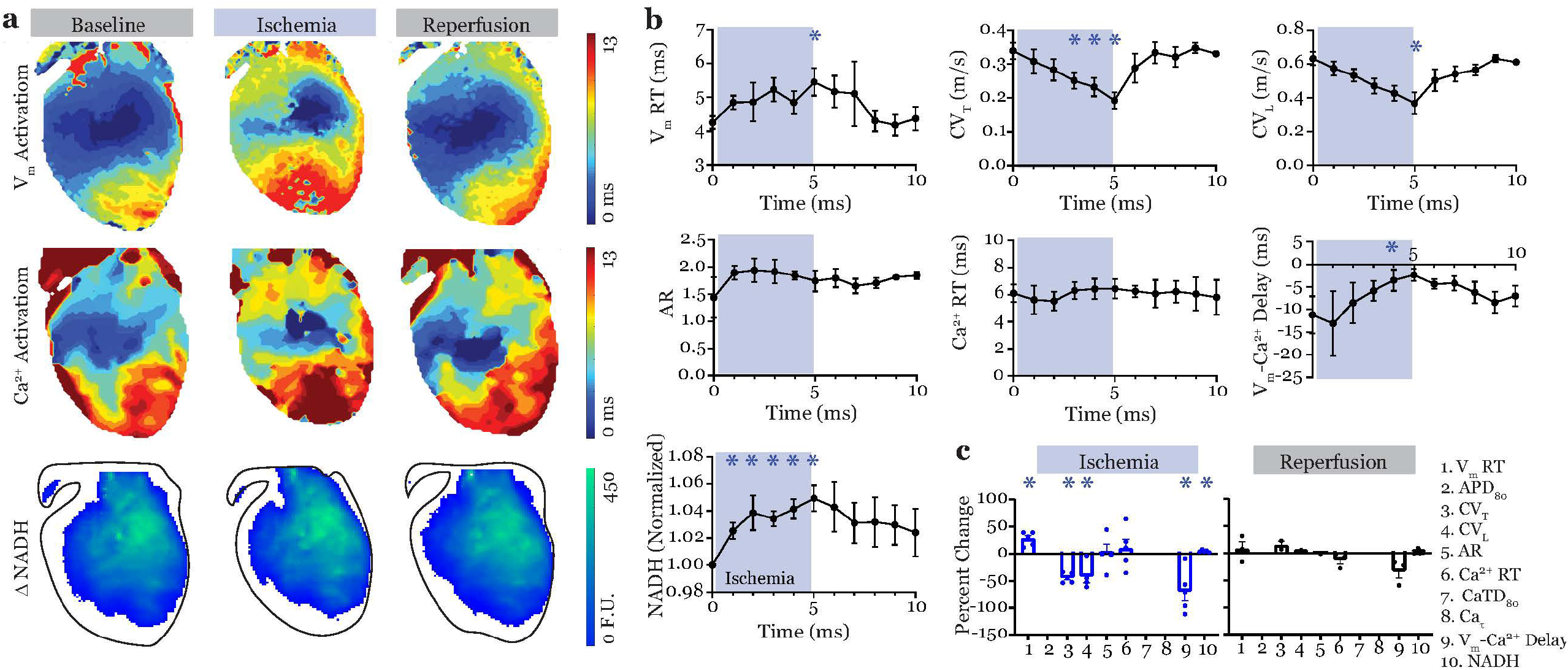
Triple parametric optical mapping in cardiac disease assessment. **a)** Representative V_m_ activation maps (top), Ca^2+^ activation maps (middle) and NADH intensity maps (bottom) recorded from the same mouse heart during baseline, ischemia and reperfusion conditions. All maps in a given column were obtained by simultaneous imaging. **b)** Summary restitution properties of seven parameters measured simultaneously by triple parametric optical mapping. Three parameters – APD_80_, CaTD_80_ and Ca^2+^ decay were not measurable due to motion artifacts in the repolarization phase due to absence of electromechanical uncoupling. **c)** Ten parameter panels (only 7 parameters reported) demonstrating changes in cardiac physiology during ischemia and restoration during reperfusion. Data presented as mean ± s.e.m., n=5, * denotes p<0.05 versus baseline using paired Student’s t-test with Bonferroni correction.

As expected, the first parameter that was significantly altered during ischemia was NADH intensity. A quick increase in NADH levels was observed, as early as 1 min into ischemia. This was followed by changes in electrophysiology. Specifically, ischemia slowed CV_T_ at 3 mins and then slowed CV_L_ and prolonged V_m_ RT at 5 mins. No changes in AR or Ca^2+^ RT were measured during the 5 min ischemic protocol. Lastly, V_m_-Ca^2+^ delay was also decreased by ischemia, possibly due to prolonged V_m_ upstroke but unaffected Ca^2+^ upstroke.

Reperfusion restored all tested parameters to pre-ischemic baseline values as quickly as 1 min after start of perfusion. Such a quick response is possibly due to the short duration of the preceding ischemia.

## Discussion

We present here the first application of triple parametric optical mapping for simultaneous measurements of V_m_, Ca^2+^ and NADH, which allow studying MECC. We demonstrated the significance of this methodology in drug testing and cardiac disease studies by performing triple-parametric optical mapping in mouse hearts during blebbistatin, 4-AP and verapamil treatments as well as during ischemia and reperfusion. We report that while blebbistatin and 4-AP modulated multiple parameters of cardiac physiology, the effects of verapamil were focused to a single parameter. Specifically, verapamil treatment induced prolongation of Ca^2+^ RT which could be expected with an I_CaL_ blocker. On the other hand, blebbistatin prolonged V_m_ and Ca^2+^ RTs while 4-AP caused prolongation of APD, Ca^2+^ RT as well as slowing of CV_T_. This methodology was also applied to investigate the acute effects of ischemia and reperfusion. While ischemia affected multiple parameters including increase in V_m_ RT and NADH as well decrease in CV_T_, CV_L_ and V_m_-Ca^2+^ delay, reperfusion restored all these parameters to baseline values. By simultaneously measuring multiple aspects of cardiac function, we determined that changes in the metabolic state precedes the electrophysiological modulation during ischemia. Thus, by implementing triple parametric optical mapping, we determined unexpected off targets effects of drugs and sequence of modulation of cardiac physiology in disease.

### Effects of Blebbistatin on cardiac physiology

Blebbistatin is a selective inhibitor of myosin II isoforms found in skeletal muscles with little to no effect on other myosin isoforms. blebbistatin binds to the myosin-ADP-Pi complex and interferes with the phosphate release process, leaving the myosin detached from actin thereby arresting cellular contraction and preventing energy consumption by contraction thus reducing metabolic demand^13^. Blebbistatin is widely used as an electromechanical uncoupler to study cardiac physiology by optical methods which require arresting the heart to prevent motion-induced artifacts. Blebbistatin is more advantageous to previously used electromechanical uncouplers in that its effects on cardiac physiology are minimal^14,15^. Blebbistatin does not alter calcium transient amplitude, rise time or decay as well as effective refractory period and ECG parameters^14^. However, mixed reports on its effects on rabbit APD have been previously published with groups demonstrating that blebbistatin either does not alter APD or that it prolongs APD^14,16,17^. Differences in methodologies including experimental conditions, poor perfusion, and motion correction algorithms applied could account for some of these differences in results. For example, blebbistatin applied to an ischemic preparation is likely to reverse ischemia induced APD shortening, appearing to prolong APD.

Although the effects of blebbistatin are well studied, these studies were mostly performed on rat or rabbit hearts and tested a concentration range of 0.1 – 10 μM which is less than recently reported concentrations used in mouse hearts^18^. In this study, we used 15 μM blebbistatin to arrest heart motion during optical mapping and determined the effects of this concentration of blebbistatin on mouse cardiac physiology. Blebbistatin altered 2 of the 10 parameters measured in this study at a “normal” pacing rate (150 ms BCL, 400 bpm) for ex vivo hearts. Upstroke rise time of action potentials and calcium transients were prolonged during blebbistatin treatment. Additionally, at slower heart rate (200 ms BCL), CV_T_ was faster and this correlated with reduced NADH autofluorescence intensity. Lower NADH values could correspond to increased ATP availability which could in turn modulate the ion channel and gap junction activity that affect cardiac conduction. This example illustrates how the use of triple parametric optical mapping can uncover such interrelated MECC that support cardiac physiology.

Blebbistatin is fluorescent with solvent-specific spectral properties which could interfere with the measurement of relative changes in NADH autofluorescence intensity between treatments. Blebbistatin dissolved in DMSO (solvent used in this study) has an excitation/emission peak of 420/560 nm and the majority of the emission is above 500 nm^19^. The design of this triple-parametric optical mapping system which filters NADH optical signals using a 450±50 nm filter, thus avoids the addition of blebbistatin fluorescence in NADH signals. Furthermore, exposure of blebbistatin-perfused hearts to UV light, at intensities and durations required for NADH imaging, does not cause cytotoxicity or significant changes in its electromechanical uncoupling properties^19^.

### Effects of 4-AP on cardiac physiology

4-AP is a potent inhibitor of the I_to_ currents and is used in treatment of multiple sclerosis. Inhibition of I_to_, a Phase 1 repolarizing current could cause prolongation of APD. In cardiac tissue, 4-AP has been demonstrated to have a biphasic effect where APD shortening^20^ is observed at lower concentrations but APD prolongation is induced at higher concentrations (> ~5 mM)^21–23^. This could be due to the inhibitory effects of 4-AP on other ion currents like I_Kur_ and hERG^22,24^. In the presence of isoproterenol, 4-AP also promotes EADs and DADs in cardiomyocytes^25^.

In this study, we observed APD prolongation by 7 mM 4-AP treatment, as expected, at all studied pacing rates. Additionally, this APD prolongation prevented 1:1 capture at pacing at rates faster than 125 ms BCL in mouse hearts. However, we report here that the effects of 4-AP on cardiac physiology extend beyond the expected blocking of the I_to_ current.

An inverse relationship between Phase 1 repolarization and calcium transient amplitude has been reported^26^. Decrease in Phase 1 repolarization rate has been demonstrated to increase I_CaL_, calcium transient amplitude and rise time. In our study, we also report an increase in the calcium transient rise time in hearts treated with 4-AP further supporting this relationship. However, the non-ratiometric calcium dyes used in this study did not allow the accurate quantification of calcium transient amplitude.

Lastly, we also demonstrated that 4-AP slows CV_T_ in mouse hearts. Although the effects of 4-AP on cardiac conduction has not been previously reported, it has been demonstrated to restore conduction in injured neurons^27,28^. Although, we did not test the effects of 4-AP in the context of injury, we report that 7 mM 4-AP reduces conduction velocity in mouse hearts. The dose dependence and underlying mechanism of this response will need further investigation.

### Effects of Verapamil on cardiac physiology

Verapamil, an I_CaL_ blocker used to treat angina, hypertension, tachycardia and other cardiac diseases also has hERG channel blocking properties^29,30^. It is probably due to its inhibitory effect on both potassium and calcium currents that the APD response to verapamil treatment has produced mixed results in previous studies. Verapamil-induced APD prolongation, shortening and no change have been previously reported^31–33^. The varying dose-dependent effects of verapamil on I_CaL_ versus hERG current as well as differences in experimental models and tissues could explain some of these differing results. Verapamil has also been reported to have age-dependent effects on cardiac electrophysiology^34^. In this study, we report no change in APD in mouse hearts treated with 1 μM verapamil.

Furthermore, verapamil does not alter depolarization-related parameters. Verapamil has been reported to not alter the rate of depolarization in cells with sodium-dependent depolarization^35^ or alter conduction velocity^36^. In line with these findings, we report no changes in V_m_ RT and CV in mouse hearts treated with verapamil.

The effects verapamil on calcium handling are many. Inhibition of I_CaL_ by verapamil has been shown to reduce the amplitude of calcium transients and contractility^31,37^. It has also been demonstrated to suppress calcium transient alternans and reduce spontaneous calcium release^38,39^. We report here prolonged Ca^2+^ RT in verapamil-treated mouse hearts. Increase in Ca^2+^ RT despite reduced calcium transient amplitude, could suggest significantly decreased I_CaL_ and calcium release from the sarcoplasmic reticulum. Lastly, we also report here that verapamil did not induce any changes in calcium reuptake as indicated by no changes in CaTD and Ca_τ_ parameters.

### Modulation of cardiac physiology by Ischemia/Reperfusion

Ischemia is a condition which is caused by the reduction or lack of blood supply to heart tissue. Ischemia modulates multiple parameters of cardiac physiology, including all three components of physiology measured in this study. Although the acute and chronic effects of ischemia are well-established, this study is the first to simultaneously assess cardiac electrical, calcium handling and metabolic functions to determine the complex sequence of MECC. Some of the well-known effects of acute ischemia include ATP reduction (NADH increase), APD shortening, V_m_ RT increase, CV slowing, calcium alternans and spontaneous calcium release^2,9,10,12^.

Changes in NADH due to acute cardiac ischemia occur within 15 s and reperfusion can restore it to baseline within 60 s^40^. In this study, we used a 5 min no flow ischemia model to measure the changes in cardiac physiology during acute ischemia and reperfusion. Ischemia increased NADH levels in the tissue at the earliest time point measured (1 min) and remained elevated throughout the ischemic period. This was the first of the ten parameters measured to be modulated suggesting that ATP depletion underlies most other physiological effects of ischemia. Next, at 3 mins of ischemia, CV_T_ slowing was observed. This was followed by decrease in CV_L_ and and increase in V_m_ RT. The effect of reduced ATP on the phosphorylation state of depolarizing sodium current and gap junctions could underlie these effects. It is also important to note that the effects of ischemia on V_m_ RT could be underestimated because the optical action potential recorded in each pixel is an average of multiple cardiomyocytes. Therefore, it is possible that V_m_ RT is increased sooner in the ischemic period than measured with this approach. Lastly, ischemia also reduced the V_m_-Ca^2+^ delay possibly due to slower depolarization (increased V_m_ RT).

Reperfusion restored all measured parameters to pre-ischemic values within 1 min of restarting the perfusion to the heart. The short duration of the ischemic period could account for the immediate return to baseline conditions. Future studies aimed at determining the sequence of restoration of cardiac physiology during reperfusion could include prolonged ischemia periods or more frequent recordings during the reperfusion period.

## Conclusions

This study demonstrates for the first time the application of triple-parametric optical mapping, which allows studying metabolism-excitation-contraction coupling in the heart. Here, we applied this methodology for drug cardiotoxicity testing and to study the modulation of cardiac physiology during ischemia/reperfusion. We identified ten parameters of cardiac physiology related to electrical excitation, calcium handling and metabolism that give important information on the state of the heart. We developed a ten-parameter panel (TPP) graph which can give a quick overview of the effects of drugs or diseases on the heart. Using this approach, we determined the effects of blebbistatin, 4-AP, and verapamil on mouse cardiac physiology. While blebbistatin and 4-AP altered multiple aspects of cardiac physiology, the effects of verapamil were limited to calcium transient upstroke as expected with a calcium channel blocker. This demonstrates that triple parametric optical mapping is a valuable tool to study cardiotoxicity of drugs in preclinical trials, particularly to identify off-target effects. Current drug testing is limited primarily to QT interval testing. This field could greatly benefit from a more comprehensive assessment of cardiac physiology as is the case with triple parametric optical mapping. Lastly, we also applied this methodology to determine the sequence of modulation of the multiple facets of cardiac physiology during acute ischemia. Simultaneously measuring the three facets of cardiac physiology identified that changes in metabolism during acute ischemia precede the effects on electrophysiology. The critical applications of this methodology demonstrate the need and the significance of triple parametric optical mapping.

### Limitations

The effects of ischemia on repolarization/calcium reuptake parameters were not measurable because optical mapping was only performed in the absence of the electromechanical uncoupler blebbistatin. This approach was implemented because the changes in NADH due to ischemia were not accurate in the electromechanically uncoupled hearts, as blebbistatin reduces metabolic demand.

Another limitation of blebbistatin is its fluorescence overlapping with NADH, which requires careful consideration of the data recorded from blebbistatin-treated heart in metabolically compromised states.

Hearts were only subjected to 5 mins of ischemia because beyond this time point, the quality of the optical signals deteriorated and did not allow appropriate analysis. Future studies will explore newer dyes with improved signal quality even during ischemia.

## Online Methods

All experimental protocols were approved by the Institutional Animal Care and Use Committee at The George Washington University and are in accordance with the National Institutes of Health Guide for the Care and Use of Laboratory Animals.

### Triple Parametric Optical Mapping system set up and alignment

The triple parametric optical mapping system uses three CMOS cameras (MiCAM05, SciMedia) focused on the same field of view through a tandem lens configuration as illustrated in Figure 1A. All filter cubes, mechanical and stage components of the system were custom 3D printed and were previously published for dual parameter optical mapping. Two modifications were made to these parts to accommodate an extra camera in this current system (Figure 1b). Open-source files of these components are also available at Github (https://github.com/optocardiography).

Two excitation light sources at 365 nm and 520±5 nm were used in episcopic illumination mode to excite the dyes or induce autofluorescence in the cardiac tissue. The emitted light is collected by a 1X objective lens (SciMedia) which focuses the light at infinity. This infinity correction allows multiple cameras to be introduced in the light path. Light at different wavelengths were separated using dichroic mirrors and filters as illustrated in Figure 1c and passed through a second lens (projection lens) before being recorded using the CMOS cameras. First, light below 510 nm is split from the straight light path by a dichroic mirror which is then filtered by a 450±50 nm filter and directed to the first CMOS camera to record NADH autofluorescence. Next, Rhod2-AM (intracellular calcium) signal is split from the straight light path using a 630 nm dichroic mirror, filtered using a 590±33 nm filter and recorded using the second CMOS camera. Finally, the RH237 (transmembrane potential) signals at wavelengths above 630 nm are filtered through a 690±50 nm filter and recorded using the third CMOS camera.

At the start of each experiment, all three cameras were focused and aligned. Each camera was attached to its own projection lens and focused at infinity. The projection lens/camera unit was then attached to the filter cubes. A focusing target was positioned in front of the objective lens and the position of the target was adjusted until all three cameras were in focus. Next, the three cameras were spatially aligned using the Camera Calibration function in the BVWorkbench software (Brainvision) and by adjusting the angle of the dichroic mirrors. Once, all cameras were focused and aligned the system is ready for use.

### Langendorff perfusion

Adult male and female mice on C57BL/6 background were used in this study. Mice were anesthetized by isoflurane inhalation and cervical dislocation was performed. Hearts were quickly excised following thoracotomy and the aorta was cannulated. The heart was attached to a Langendorff perfusion system, hung in a vertical position in a temperature-controlled bath and perfused with a modified Tyrode’s solution containing (in mM) 130 NaCl, 24 NaHCO3, 1.2 NaH2PO4, 1 MgCl2, 5.6 Glucose, 4 KCl, 1.8 CaCl2, pH 7.40 and bubbled with carbogen (95% O2 and 5% CO2) at 37°C. Perfusion pressure was maintained at ~80 mmHg by adjusting the flow rate between 1 and 1.5 ml/min. A platinum bipolar electrode was placed at the center of the anterior surface residing the middle of the field of view. Gentle pressure was applied to the back of the heart as it was pushed up to the front optical glass of the bath using a paddle, allowing the pacing electrode to be held in place. Electrical stimuli were applied to determine the threshold of pacing. Hearts were paced at 1.5X pacing threshold amplitude and 2 ms stimulus duration, only during optical recording. Hearts were paced at varying rates as listed in the figures to determine the restitution properties of the heart under each condition.

### Optical Mapping

The heart was subjected to a 10 min equilibration period followed by staining with the voltage- and calcium-sensitive dyes. A 1 ml mixture of RH237 (30 μl of 1.25 mg/ml dye stock solution + 970 μl Tyrode’s solution) was prepared and immediately injected into the dye port above the cannula over a 3-5 min period. This was followed by a 5 min washout period. Similarly, a 1 ml mixture of Rhod2-AM (30 ul of 1 mg/ml dye stock solution + 30 μl Pluronic F-127 + 940 μl Tyrode’s solution) was prepared and immediately injected into the dye port over a 3-5 min period. This was followed by 5 min dye washout period.

The heart was then illuminated by two LED excitation light sources at 365 nm and 520±5 nm wavelengths. While the former induces autofluorescence of NADH in the tissue, the latter excites both RH237 and Rhod2-AM dyes. Control optical recordings of NADH, V_m_ and Ca^2+^ were simultaneously acquired at 1 kHz sampling rate.

### Electromechanical uncoupling and drug treatment

After Control recordings, hearts were treated with 15 μM blebbistatin for 20 mins and optical recordings were acquired as above. Next, hearts were treated with 4-AP (7 mM) and verapamil (1 μM) one at a time, in the presence of blebbistatin. Optical recordings were once again acquired for each of these conditions as described above.

### Ischemia

In a separate set of hearts, after the initial equilibration period and dye staining/washout, hearts were subjected to 5 mins of no flow ischemia followed by 5 mins of reperfusion. Hearts were continually paced at 300 ms BCL throughout this period. Optical recordings were acquired prior to (baseline), during (ischemia) and after (reperfusion) this period of no perfusion at 1 min intervals. Ischemia period was limited to 5 mins because beyond this point, the optical signals were of very poor quality which would not allow suitable analysis.

### Data Analysis

Optical data of NADH, transmembrane potential and calcium were analyzed using a custom Matlab software, Rhythm 3.0, which is available in an open-source format on Github. Rhythm 3.0 which is an upgraded version of Rhythm 1.2, incorporates NADH visualization and analysis features as well as the ability to calculate delay in transmembrane potential to intracellular calcium activation (V_m_-Ca^2+^ Delay).

In this study, 10 different parameters were measured from the 3 optical signals that were simultaneously recorded under each condition. These include, from transmembrane potential: (1) rise time (V_m_ RT), (2) action potential duration (APD_80_), (3) transverse and (4) longitudinal conduction velocity (CV_T_ and CV_L_) and (5) anisotropic ratio (AR), from intracellular calcium: (6) rise time (Ca^2+^ RT), (7) calcium transient duration (CaTD_80_), (8) calcium decay time constant (τ), (9) V_m_-Ca^2+^ delay and (10) NADH fluorescence intensity. The rate-dependence of each of these parameters is indicated in the parameter vs BCL graphs while the summary of effects of each condition is illustrated in the TPP graphs. The values reported in the TPP graphs correspond to BCL = 150 ms which is considered “normal” heart rate (400 bpm) for an ex vivo mouse heart preparations.

Upstroke rise time (RT) from V_m_ and Ca^2+^ signals were measured as the time period from 20 to 90% of the upstroke of the action potential and calcium transient, respectively. APD_80_ and CaTD_80_ were measured as the time interval between activation time (time of maximum first derivative of the upstroke) and 80% of repolarization and calcium transient decay, respectively. CV_L_ and CV_T_ in the parallel and perpendicular direction to fiber orientation, respectively, were calculated using differences in activation times and known interpixel distances. Anisotropic ratio was calculated as the ratio of CV_L_ to CV_T_. Calcium decay time constant (τ) was determined by fitting an exponential to the last 50% (50 – 100%) of the calcium transient decay phase. V_m_-Ca^2+^ delay was defined as the time interval between the activation time of the V_m_ signal minus the activation time of the Ca^2+^ signal. Lastly, NADH was measured as the average of the absolute autofluorescence intensity value in the optical recording. NADH intensity in each heart was normalized to Control.

### Statistics

All data are reported as mean ± standard error of the mean (SEM). One-way ANOVA tests were performed to determine significant differences between groups and paired, two-tailed Student’s t-tests were performed for post hoc analysis. Bonferroni correction was applied for multiple comparisons.

## Acknowledgements

This work was funded by Leducq Foundation (project RHYTHM), American Heart Association SCD SFRN grant, and NIH grant R44 HL139248 to IRE and an American Heart Association Postdoctoral Fellowship (19POST34370122) to SAG.

## Author contributions

Study conception: SAG, IRE; Study design: SAG; Data acquisition, analysis and interpretation: SAG; Designing and 3D printing hardware: SAG, ZL; Preparing analysis software: SAG; Manuscript drafting and figure preparation: SAG; Revision and final version approval: SAG, ZL, IRE.

